# FIRST REPORT OF HALOBACTERIA DOMINANCE IN A TROPICAL CAVE MICROBIOME

**DOI:** 10.1101/2021.12.02.470950

**Authors:** Caio César Pires de Paula, Dagmara Sirová, Hugo Sarmento, Camila Cesario Fernandes, Luciano Takeshi Kishi, Maria Elina Bichuette, Mirna Helena Regali Seleghim

## Abstract

Scarce studies on microbial diversity in tropical caves have been published, a subterranean system still neglected from a microbiological point of view. Although most published studies are about temperate caves, usually archaeas and fungi have less attention than bacterial communities. Here, the microbiome structure and composition in a tropical cave system, as well the main environmental drivers, were studied during the wet and dry season. Soil and sediments from three different habitats at the cave (surface, entrance cave and dark zone) were sampled. Samples were characterized (temperature, air and substrate humidity, salinity, pH, nitrogen and organic carbon content, and chemical composition) and the microbiome was assessed by high-throughput sequencing, using amplicon sequencing (16S and ITS). Prokaryotic communities were dominated by *Halobacteria, Actinobacteria* and *Bacilli*, while fungal communities showed high abundance of *Sordariomycetes*. Microbiomes from the cave entrance, where a significantly elevated salinity levels were found, supported up to 63% of *Haloarchaea* compared to the other habitats studied. Differences in community structure were significant between habitats, but no influence of the season was observed. Main environmental drivers of community assembly included nitrogen and organic carbon content, temperature, and salinity. This is the first report of *Halobacteria* dominance in cave habitats, where they likely play important roles in nitrogen and phosphorus cycles. The cave entrance had lower diversity, but higher degree of microbial endemism, which characterize it as an important cave ecotone. The prevalence of heterotrophic microbial groups implies trophic structure based on detritivores, particularly in the dark zones. Our study brings new insights on microbiome composition in the underexplored tropical cave habitats.

## 1 INTRODUCTION

Caves are generally classified as extreme environments due to the prevailing oligotrophy and lack of light to support photosynthesis. They therefore represent highly specialized ecosystems with no autochthonous photosynthesis-based primary production, dependent mostly on allochthonous carbon inputs [1]. Microbial biodiversity, function, and community dynamics in caves are still considered somewhat of a “black box”. Although the number of published results on cave microbiology has increased over the past decade, most of them were based on temperate cave habitats, and the information on tropical caves are still scarce [2]. Tropical caves are in regions differing significantly in environmental conditions regarding the temperate caves, which include, in general, climate with higher temperatures, a dense tree canopy whole year, due the lack of a dormant period in the winter or dry season, and higher primary production [3]. Such differences can influence the structure and dynamics of microbial communities in caves [4 – 5], highlighting the importance to understand the microbial ecology in tropical cave environments over a seasonal cycle.

Even though the published literature covers a wide variety of microbial habitats within the caves such as soils, sediments, stream water, and rock surfaces, most information is limited to describing the *Bacteria* domain only. Indeed, caves show high abundances of *Proteobacteria*, presumably involved in nitrogen fixation, along with significant populations of *Actinobacteria*, with a suggested role in carbon turnover [6]. Nevertheless, studies evaluating the presence and dynamics of other microbial groups in the caves - the *Archaea* and *Eukarya* – are rare. Recently, after a microbiome definition update, researchers have highlighted the importance of considering all microorganisms belonging to different kingdoms (prokaryotes: bacteria, archaea, and micro-eukaryotes: as fungi), as well as their functions and interactions, as part of microbiome studies [7].

Next generation sequencing technology advances in the last decade made it easier to assess the uncultured microbial diversity in cave environments and recent studies have demonstrated the presence of *Archaea*, most frequently from the phyla *Thaumarchaeota* and *Euryarchaeota* [8 – 10]. Methanogenic archaea (MA) and ammonia-oxidizing archaea (AOA) are the best-characterized archaeal groups and these microorganisms are targeted in studies of subterranean environments due to their ecological importance in biogeochemical cycling [11]. These microorganisms are involved in the terminal steps of carbon flows, nitrification processes [12], primary production [13], and denitrification [14] in terrestrial and aquatic environments. There are currently no *Thaumarchaeota* in culture. Although these are the most frequently encountered *Archaea* in caves, we have very limited information on the possible biogeochemical and ecological roles of *Thaumarchaeota* in these specific environments [11, 15 - 16]. Most of what we do know has been inferred from metagenomics studies: *Thaumarchaeota* are likely to play an important role in ammonia oxidation in temperate caves [10]. Some cave microbiomes contain a large proportion of *Euryarchaeota*. For instance, four *Euryarchaeota* classes - *Methanomicrobia, Thermoplasmata, Halobacteria* and *Methanobacteria* - were found in Indian caves systems [8]. The *Euryarchaeota* include extreme halophiles, sulfate reducers, thermophilic heterotrophs, and methanogens [17].

More than 1150 species of fungi, belonging to 550 genera, have already been reported in caves worldwide [18]. The most identified fungi in caves belongs to the phyla Ascomycota, Basidiomycota, and Zygomycota [19]; however, the relevance of these findings should be viewed with caution, as these studies utilize cultivation-dependent techniques. The difficulty of laboratory cultivation, especially those fungi colonizing rocks or low metabolic activity, limits the knowledge of the real diversity, as well as the exploration of the ecological roles performed by these microorganisms in the cave systems. Cave fungi are decomposers or parasites, although they can also aid in speleogenesis processes through the precipitation of secondary minerals [20]. Fungi are the main saprophytic organisms in cave systems and play an important role in the food web [19]. However, pathogenic fungi receive more attention in cave studies due to health and economic concerns, which mainly include *Histoplasma capsulatum* and *Pseudogymnoascus destructans*.

Taking into account i) the tropical caves are neglected from a microbiological point of view and ii) the gap in the information about achaeas and fungi in cave microbiomes, the main goal of the present study were the analysis the structure and composition of the microbiomes (here including archaea, bacteria and fungi), in different habitats within a tropical cave system, to evaluate the main environmental factors that drive the structure of the microbiomes, and to discuss the possible role of the cave microorganisms in the trophic structure of the subterranean habitats. This is the first research in Brazilian caves using new generation sequencing to assess fungal and prokaryotic diversity. Belonging to the several larger karst areas in Brazil, Terra Ronca State Park (PETeR) is a state conservation unit located in Central Brazil. PETeR harbor the main complex of cave systems in Brazil, including several superficial and subterranean drainages, with great potential for the transport of organic matter, causing accumulations of debris in some caves. This causes high habitat differentiation, accompanied by high richness of subterranean terrestrial and aquatic taxa [21 – 25]. Although the biology of these biodiversity hotspots, especially that of the subterranean fauna, is well known, no studies focused on the microbial assemblages were published so far.

## 2 MATERILAS AND METHODS

### 2.1 STUDY SITE

The Terra Ronca State Park (PETeR) (46°100’- 46°300’S; 13°300’ – 13°500’ W), located within São Domingos city (Goiás State, central Brazil), has a large subterranean system formed by rivers arriving from the Serra Geral Plateau, a morphological feature that originated in the Cretaceous, in the sandstones of the Urucuia Formation [26]. The PETeR is a karst area crossed by parallel streams running westwards to join the Paraná River, a tributary of the Upper Tocantins River, in the Amazonas Basin. The study area belongs to the Cerrado phytogeographical domain (a savanna-like vegetation). The climate is tropical semi-humid with a mean annual precipitation of about 1270 mm yr^−1^ [27]. The wet season extends from November to April, with rainfall essentially absent between May and October (dry season). Although the area is a Conservation Unit, the park is threatened by anthropogenic impact, such as deforestation for agriculture and uncontrolled tourism, and the springs of the main rivers are outside the boundary of the Conservation Unit, a place used for cattle, crops, consequently silting and polluting rivers and affecting the cave systems [28].

The Lapa da Terra Ronca I cave (TR cave) is a part of the Terra Ronca-Malhada cave system (Fig 1, Fig S1). The cave has an entrance of ca 100 meters high and 120 meters wide, with a built altar measuring 760 meters long and 100 meters high where the local religious ceremony of “Bom Jesus da Lapa” takes place at the beginning of August.

**Fig 1.**
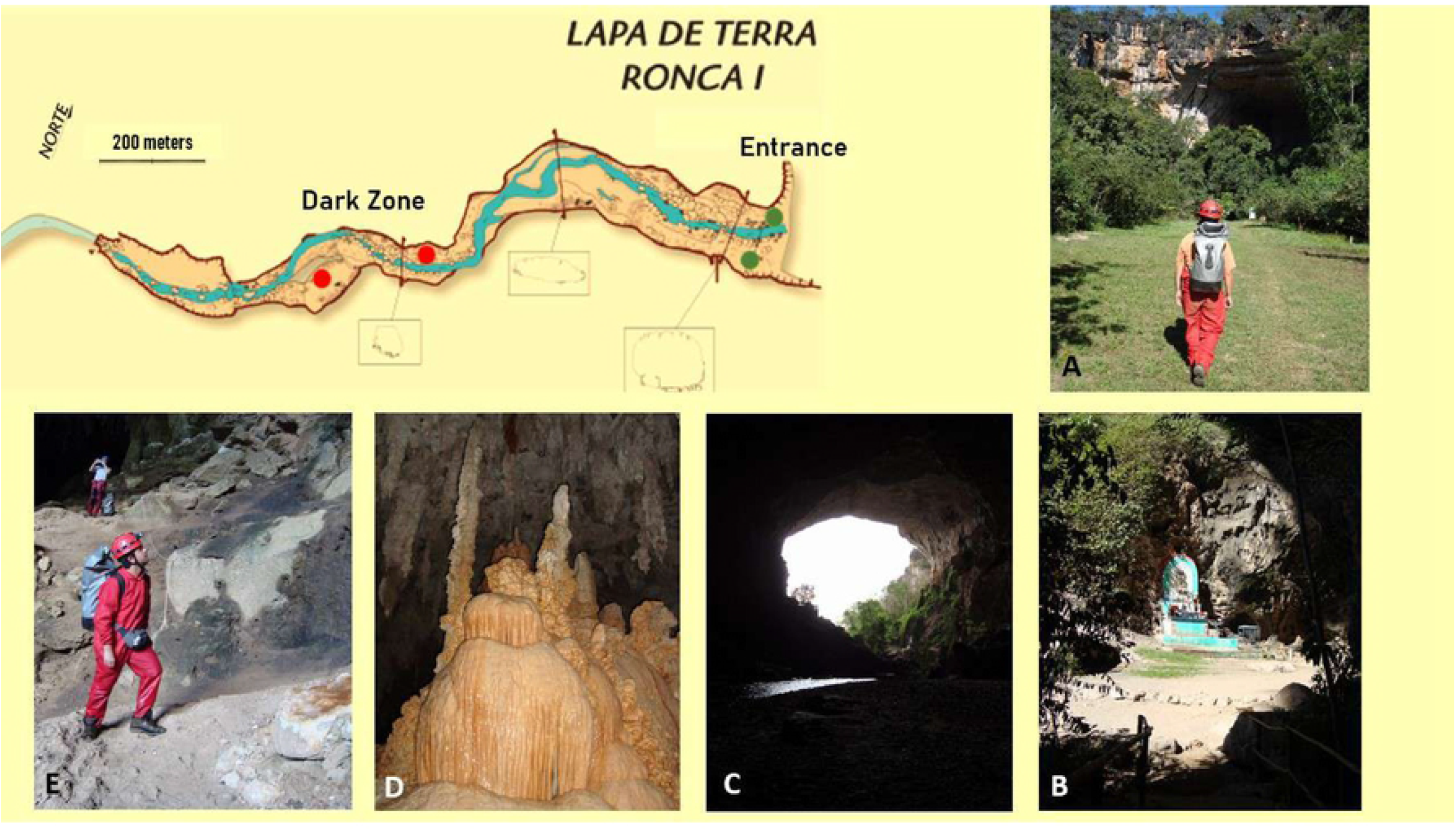
Map of Lapa de Terra Ronca I cave (TR cave) located in a Brazil Central region, and the photos shows the surface (A), external (B) and internal (C) view of entrance area and the dark zone (D and E). The dots show the replicates samples in the map at entrance cave (green) and dark zone (red). Surface samples were collected 50 meters far from cave entrance (map: Grupo Bambuí de Pesquisas Espeleológicas – GBPE).

### 2.2 SAMPLING

Samplings were conducted in April 2016 (wet season) and October 2016 (dry season) (license n° 28992-11/ICMBio/SISBIO and n° 14886/2010/Secima, Goiás). One square of approximately 0.25 m^2^ was sampled in three distinct habitats of the cave: surface (cave exterior – around 50 meters far from cave entrance), cave entrance and the dark zone (no sunlight). All the different habitats were sampled in replicates. Approximately 300 g of substrate (soil or cave sediment, up to 10 cm in depth) were collected at five different points of the square area at each site forming a composite sample. Samples were collected with the aid of a shovel and stored in sterile plastic bags. The samples were transported to the laboratory in coolers, homogenized, sieved (2 mm mesh) and stored in the refrigerator at 4 °C.

### 2.3 PHYSICAL AND CHEMICAL PARAMETERS

Temperature (°C), air humidity (%) and luminosity (Lux) were measured at each sampled area with a minimum time interval of 1 minute between measurements (Thermo-hygrometer Instruntherm THAL-300, 0.1 resolution and ± 5.0% accuracy). Substrate pH was measured at the substrate:water ratio of 1:2.5 (weight/weight) and substrate salinity was estimated using a refractometer. The moisture in the cave sediment and soil samples was estimated by the gravimetric method with drying at 105 °C for 20 h to 7 days after sampling, and the results expressed as dry weight percentage. Organic carbon (OC) concentrations were measured colorimetrically using the method of [29]. Total nitrogen content (TN) was determined by Kjeldahl digestion followed by ammonia distillation (indophenol blue method) [30]. The composition of the subterranean substrate was assessed by scanning electron microscopy (SEM), together with chemical analysis by energy dispersive spectroscopy (EDS). An Oxford EDS coupled to a FEI Quanta 250 SEM was used to examine the chemical composition of the samples. The substrate was adhered to a double-sided copper tape mounted on an aluminum stub for observation [31]. Fifteen ESEM images and corresponding EDS spectra of elements were acquired for each sample on average.

### 2.4 DNA EXTRACTION AND SEQUENCING

DNA was extracted from 0.25 g of each homogenized sample using the MoBio PowerSoil DNA extraction kit (MoBio Laboratories, Carlsbad, CA, USA) following manufacturer’s instructions. The quality and quantity of extracted DNA were verified by the examining products on TBE agarose gels and by measuring the ratio of absorbance at 260 and

280 nm, and 260 and 230 nm, with a Thermo Scientific Nanodrop 2000c Spectrophotometer. For prokaryotes, the V3-V4 region of the 16S rDNA genes was amplified using the primer pair 341F (5′-CCTACGGGNGGCWGCAG-3′) and 805R (5′-GACTACHVGGGTATCTAATCC-3′). The ITS1 (5′-GCATCGATGAAGAACGCAGC-3′) / ITS4 (5′-TCCTCCGCTTATTGATATGC-3′) primers were used to assess the diversity of fungi [32 – 33]. Briefly, for a 25μL PCR reaction with 8.5 μl of Kapa High-fidelity HOTSTART ready MIX, 0.1 μM of each primer, 10 μl of PCR-grade water, and 10 ng of DNA extract were used. The 16S rDNA amplification conditions were 95°C for 3 min, followed by 35 cycles at 95 °C for 30 s, 55 °C for 1 min 15 s, 72°C for 45 s, and finally 72°C for 5 min. The ITS amplification had an initial stage of 95 °C for 3 min, 35 cycles of 95 °C for 15 s, 60 °C for 15 s, 72 °C for 45 s, and finally 72 °C for 10 min. PCR products were purified with the AMPURE XP magnetic bead kit (Bechman Coulter) and indexed with the Nextera XT kit V2 (Illumina) to separate samples for sequencing. A second step of purification with magnetic beads followed, and then the metagenomic pool was assembled with 5 μl of each library. High-throughput sequencing of the V3-V4 and ITS regions was performed using the Illumina MiSeq sequencing platform. ITS amplicons were sequenced at Macrogen (Macrogen Inc., Seoul, Korea), while the 16S samples at the Multi-user Laboratory of Sequencing in Large Scale and Gene Expression (São Paulo State University “Julio de Mesquita Filho”). Negative extraction and PCR controls were sequenced together with amplicons samples, and the raw sequences have been deposited in the NCBI Sequence Read Archive (SRA) under project accession number PRJNA723998 (16S) and PRJNA724003 (ITS).

### 2.5 BIOINFORMATIC ANALYSIS AND BIODIVERSITY ASSESMENT

Sequencing data were processed using UPARSE [34] in a pipeline internally implemented for 16S and ITS [35]. Paired-end reads were merged with PEAR [36]. Sequences were quality controlled with the following steps: all sequences shorter than 100 pb were discarded, followed by quality dereplication checking, OTU clustering (UPARSE algorithm, similarity ≥ 97%), and filtering of chimeras with USEARCH [37 – 38]. Taxonomic classification was performed through BLASTn using the databases SILVA 119.1 for 16S and UNITE for ITS.

All statistical analyses were carried out in R Software (R. Core Team, 2016). Environmental variables were analyzed using basic descriptive statistic (Shapiro-Wilks). Analysis of variance and Student t-tests with 5% probability threshold were also applied to verify the significance of the differences among the results. Statistical analyses of microbial community richness, alpha diversity (Shannon index) and community structure were estimated using R *microbiome* package [39]. The distance matrices of community composition were obtained using Bray-Curtis distance and, for environmental matrices, Euclidean distance was used. Relationship between environmental and microbiomes matrices were evaluated by Canonical Correspondence Analysis (CCA) using tools from a number of other R extensions, including *vegan* [40] and *ggplot2* [41]. Differences among sites sampled and the two seasons were tested using permutational multivariate analysis of variance (PERMANOVA) with Bray-Curtis distance, performing 9999 permutations using the *adonis* function [40]. Principal coordinate Analysis (PCoA) ordination using weighted Unifrac was used to evaluate the β - diversity among the sampled sites and a heatmap associated with cluster analyzes (*pheatmap* function) to identify biomarkes for each habitat [42].

## 3 RESULTS

### 3.1 PHYSICOCHEMICAL PARAMETERS

A summary of all analyzed environmental variables is presented in Table 1. The two sampling seasons differed significantly in mean temperatures and air humidity. Salinity was significantly higher at the cave entrance compared to the other sampling sites, with the difference most pronounced in the dry season (4.42%, compared to 2.36% in the wet season). The substrate in all samples were slightly alkaline (pH ranged between 7.49 and 8.88), and there was a marked trophic gradient present in both (wet and dry) sampling season: higher organic carbon content at the cave entrance (1008.20 mgC kg^-1^ and 1170.71mgC kg^-1^, respectively), followed by the surface (820.37mgC kg^-1^ and 824.48mgC kg^-1^), and the dark zone site with lowest measured organic carbon concentrations (448.93mgC kg^-1^ and 334.98 mgC kg^-1^). The surface showed higher amounts of TN in the dry season (0.075mgN kg^-1^), while higher amounts of TN at the entrance and dark zone sites were observed in the wet season (0.078 mgN kg^-1^ and 0.036 mgN kg^-1^, respectively). The detailed analysis of sample chemical composition revealed that silica (Si) was the dominant element. The cave entrance exhibited higher concentration of other essential elements, such as magnesium (Mg) and calcium (Ca) (Table 2). Samples from there were also the only ones where chlorine (Cl), sulfur (S), and phosphorus (P) were present in detectable quantities.

**Table 1.** Mean values and standard deviations of physical (substrate moisture, air temperature, air humidity and luminosity) and chemical (pH, salinity, organic carbon (OC) and nitrogen (N)) in TR cave on the surface, entrance and subterranean sample sites during wet (April 2016) and dry (October 2016) seasons.

| ENVIRONMENTAL PARAMETERS | Season | Surface | Entrance | Dark zone |
| --- | --- | --- | --- | --- |
| Substrate moisture (%) | wet | 5.60 ± 3.85<sup>a</sup> | 4.03 ± 2.54<sup>a</sup> | 3.39 ± 1.99<sup>a</sup> |
|  | dry | 1.13 ± 0.56<sup>a</sup> | 5.86 ± 1.91<sup>b</sup> | 5.91 ± 1.32<sup>b</sup> |
| Temperature (°C) | wet | 26.85 ± 0.53 | 28.33 ± 0.55 | 26.63 ± 0.98 |
|  | dry | 29.25 ± 0.10 | 29.25 ± 0.10 | 27.38 ± 0.21 |
| Air humidity (%) | wet | 73.01 ± 4.24 | 66.23 ± 0.97 | 76.71 ± 13.96 |
|  | dry | 68.86 ± 0.95 | 64.81 ± 3.93 | 68.11 ± 1.66 |
| Luminosity (Lux) | wet | 1835.33 ± 71.8 | 43.16 ± 9.3 | 0 |
|  | dry | 879.6 ± 161.6 | 233.8 ± 57.3 | 0 |
| pH | wet | 8.88 ± 0.14 | 7.49 ± 1.58 | 7.69 ± 0.25 |
|  | dry | 8.69 ± 0.29 | 7.98 ± 0.75 | 7.59 ± 0.10 |
| Salinity (%) | wet | 0.067 ± 0.007 | 2.366 ± 0.070 | 0.020 ± 0.001 |
|  | dry | 0.227 ± 0.007 | 4.426 ± 0.459 | 0.111 ± 0.014 |
| OC (mg kg<sup>-1</sup>) | wet | 820.37 ± 71.61 | 1008.20 ± 700.86 | 448.93 ± 145.67 |
|  | dry | 824.48 ± 700.84 | 1170.71 ± 1044.51 | 334.98 ± 51.23 |
| N (mg kg<sup>-1</sup>) | wet | 0.055 ± 0.040 | 0.078 ± 0.059 | 0.036 ± 0.004 |
|  | dry | 0.075 ± 0.047 | 0.025 ± 0.002 | 0.025 ± 0.007 |

**Note:** Different letters in a row means significant difference ( $p < 0.05$ ) between.

**Table 2.**
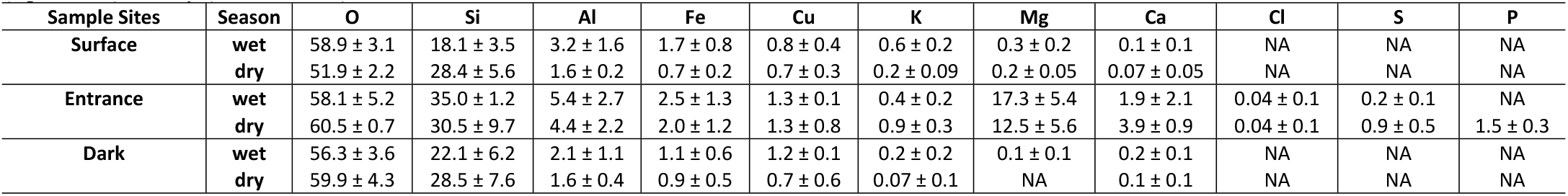
Mean values and standard deviations of chemical composition of the substrate (wt.%) in TR cave on surface, entrance and dark sites during wet (April/2016) and dry (October/2016) season.

### 3.2 MICROBIOME COMPOSITION

A total of 12 samples from three different habitats (surface, entrance, and dark zone) at TR cave were sequenced by Illumina MiSeq platform, resulting in 1,156,899 sequences for prokaryotic communities. After contig assembly, trimming, and chimera removal a total of 558,322 valid reads were obtained. Representative sequences of 3,517 OTUs were taxonomically annotated from phylum to genus levels (S2 Appendix). The a*rchaea* and *bacteria* domains showed different relative proportions at the three sampling sites. Taking into account both seasons, the entrance site showed an unusually close proportions of the two domains (44.06% *archaea* and 55.94% b*acteria*), differing significantly in this respect from the dark zone (6.62% *archaea* and 93.38% *bacteria*) and the surface (0.68% *archaea* and 99.32% *bacteria*). Particularly the cave entrance showed a larger archaea community in the dry season (75.49% *archaea* and 24.51% *bacteria*) than in the wet season (30.61% *archaea* and 69.39% *bacteria*). A total of 4 archaea phyla and 31 bacterial phyla were detected in TR cave samples. *Halobacteria* was the dominat class in archaeal communities, with *Halalkalicoccus* (54.52%) and *Halococcus* (29.46%) as the most representative taxa. *Actinobacteria* were dominant at the surface and dark zone sites, while *Bacilli* was the dominant bacterial class at the cave entrance. Fig 2A shows the main prokaryotic classes found at the sampling sites. Uncultured prokaryotic taxa comprised 24.22% at the genus level. A total of 32.5% of prokaryotic OTUs was shared between the sites, with surface showing a higher proportion of unique OTUs (16.6%), followed by the cave entrance (15.4%) and the dark zone (3.5%). About 64.4% of the prokaryotic OTUs were present in both sampling seasons.

**Fig 2.**
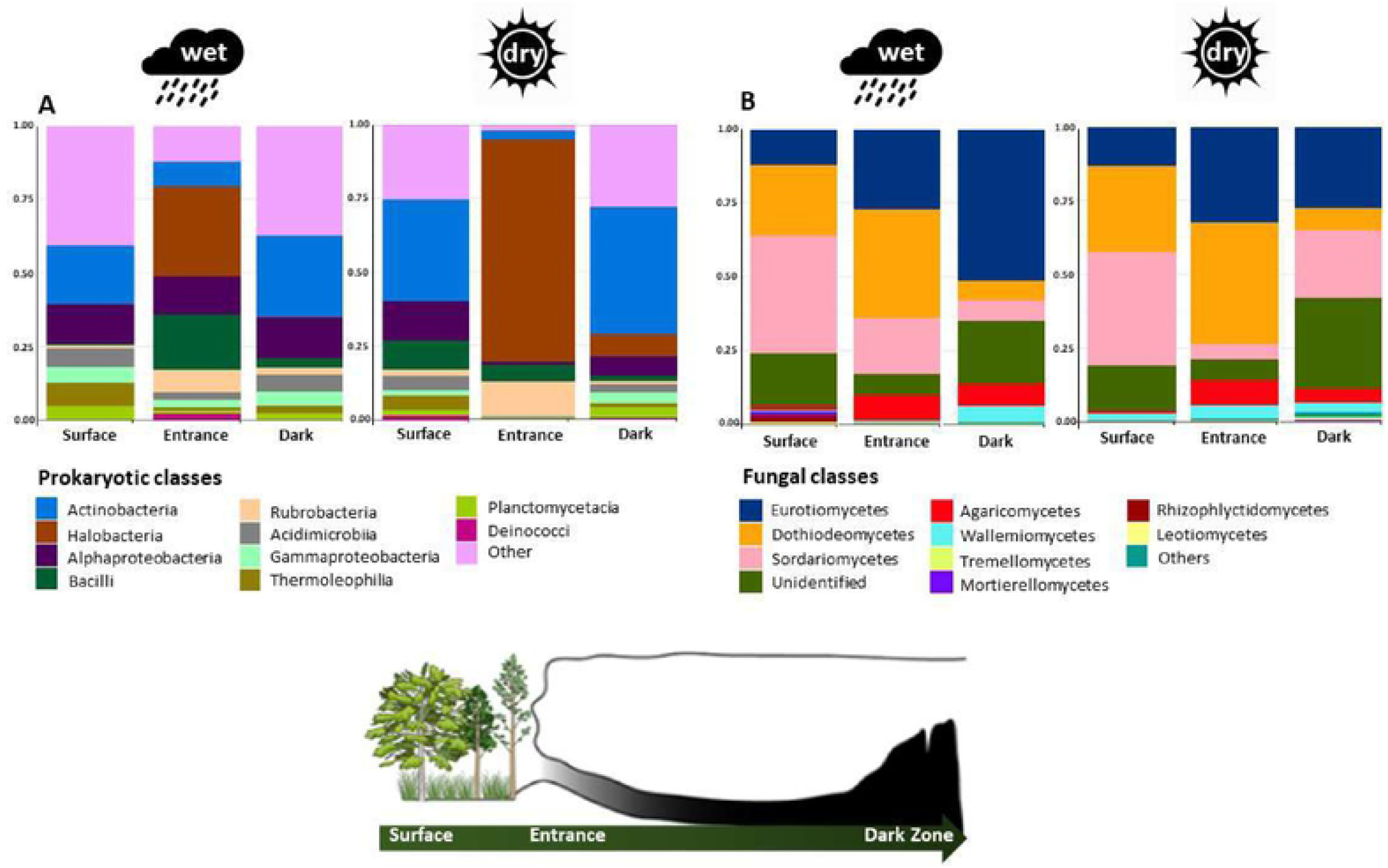
Relative abundance of prokaryotic (A) and fungal (B) communities at the Class level in TR cave during wet (April / 2016) and dry (October / 2016) season. Only the 10 most abundant classes were shown, which less abundant classes being grouped into “Others”.

Regarding fungal communities, a total of 990,619 valid reads and 1,922 OTUs were annotated from phylum to specie levels (S3 Appendix). The mean fungal OTU numbers in three different habitats ranged from 89,340 reads (surface) to 105,820 reads (entrance), with 3% cutoff. In total, 415 known genera, 193 known families, 76 known orders and 30 known classes in 7 fungal phyla were detected in the TR cave samples. *Ascomycota* was the most dominant phylum, with the relative abundance of 77.54%, followed by *Basidiomycota* (8.21%), *Chytridiomycota* (0.37%), *Mortierellomycota* (0.36%), *Mucoromycota* (0.01%), *Glomeromycota* (0.002%), and *Rozellomycota* (0.001%). The relative abundance of 10 most detected fungal taxa at the class level for all samples is showed in Fig 2B. The dataset contained fungal sequences unidentifiable at the phylum and class level (13.49% and 14.37%, respectively). *Chaetomium murorum* (17.76%) and *Aspergillus fumigatus* (15.85%) were the most abundant fungal taxa at the surface and entrance, respectively. The dominant taxon at the dark zone site was unidentified at phylum level (11.73%) and was followed by an *Aspergillus* sp. (SH186265.07FU) (6.61%). Only 3.4% OTUs were present in all sampled habitats, with higher proportion of unique OTUs at the cave entrance (42.6%) and dark zone (25.9%). About 17.1% of the fungal OTUs were present in both seasons, while 45.6% and 37.3% of the OTUs were unique in the wet and dry season samples, respectively.

### 3.3 MICROBIOME STRUCTURE AND ENVIRONMENTAL DRIVERS

The microbiome structure varied significantly among the different habitats (PERMANOVA, F2 = 1.59, p = 0.001), with no effect of the season (PERMANOVA, F2 = 0.70, p = 0.94). The same pattern can be observed analyzing only prokaryotic (PERMANOVA, F2 = 1.63, p = 0.02) or fungal (PERMANOVA, F2 = 1.34, p = 0.004) communities from the sampling sites. Based on these results, the season factor was excluded from the following analyses and the microbiome was considered as a whole, i.e. containing both prokaryotic and fungal assemblages.

Species richness (Fig 3) was highest in the surface samples (1,791.77 ± 287.24), followed by the dark zone (1,627.45 ± 1,101.80) and entrance samples (1,541.22 ± 478.32). In contrast, the Shannon index indicated that dark zones (4.38 ± 0.67) and surface (4.32 ± 0.69) had higher microbial diversity than the cave entrance (3.86 ± 0.87). Surface samples showed a slightly increased Simpson dominance (0.08 ± 0.08) compared to the entrance (0.07 ± 0.03) and dark zone (0.04 ± 0.02) samples. A cluster analysis of OTU-level diversity and PCoA based on weighted Unifrac results both showed distinct grouping of samples according to the different cave habitats, with the cave entrance cluster falling between the surface and the dark zone (Fig 4A). Overall, the core assemblage associated with each cave habitat showed a degree of specificity (heatmap analysis, Fig 5): *Cladosporium sphaerospermum* (Dothideomycetes) and *Torula* sp. (Saccharomycetes) were characteristic for surface samples, *Aspergillus* sp. (Eurotiomycetes) and *Bovista aestivalis* (Agaricomycetes) for the dark zone, while the cave entrance samples contained up to eight highly specific OTUs including *Rubrobacter* sp. (Rubrobacteria), *Halalkalicoccus tibetensis* (Halobacteria), *Cladosporium* sp. (Dothideomycetes), *Aspergillus* sp. (Eurotiomycetes), *Cladosporium halotolerans* (Dothideomycetes), *Capnodiales* sp. (Dothideomycetes), *Halalkalicoccus* sp. and *Halococcus sediminicola* (Halobacteria).

**Fig 3.**
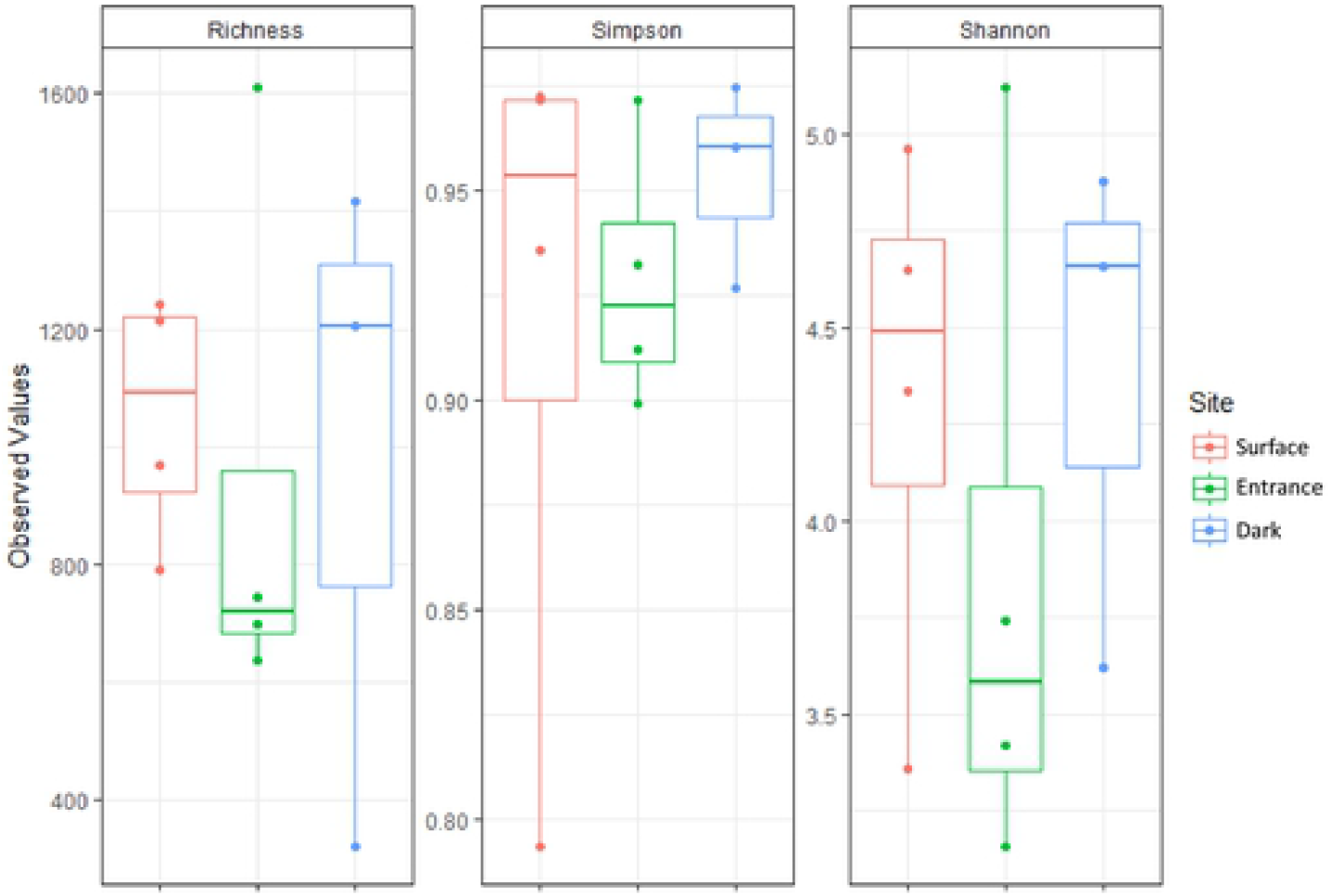
Richness (Chao1), Diversity (Shannon) and Dominance (Simpson) indices of microbial communities in three different habitats (surface, entrance and dark zone) at TR cave.

**Fig 4.**
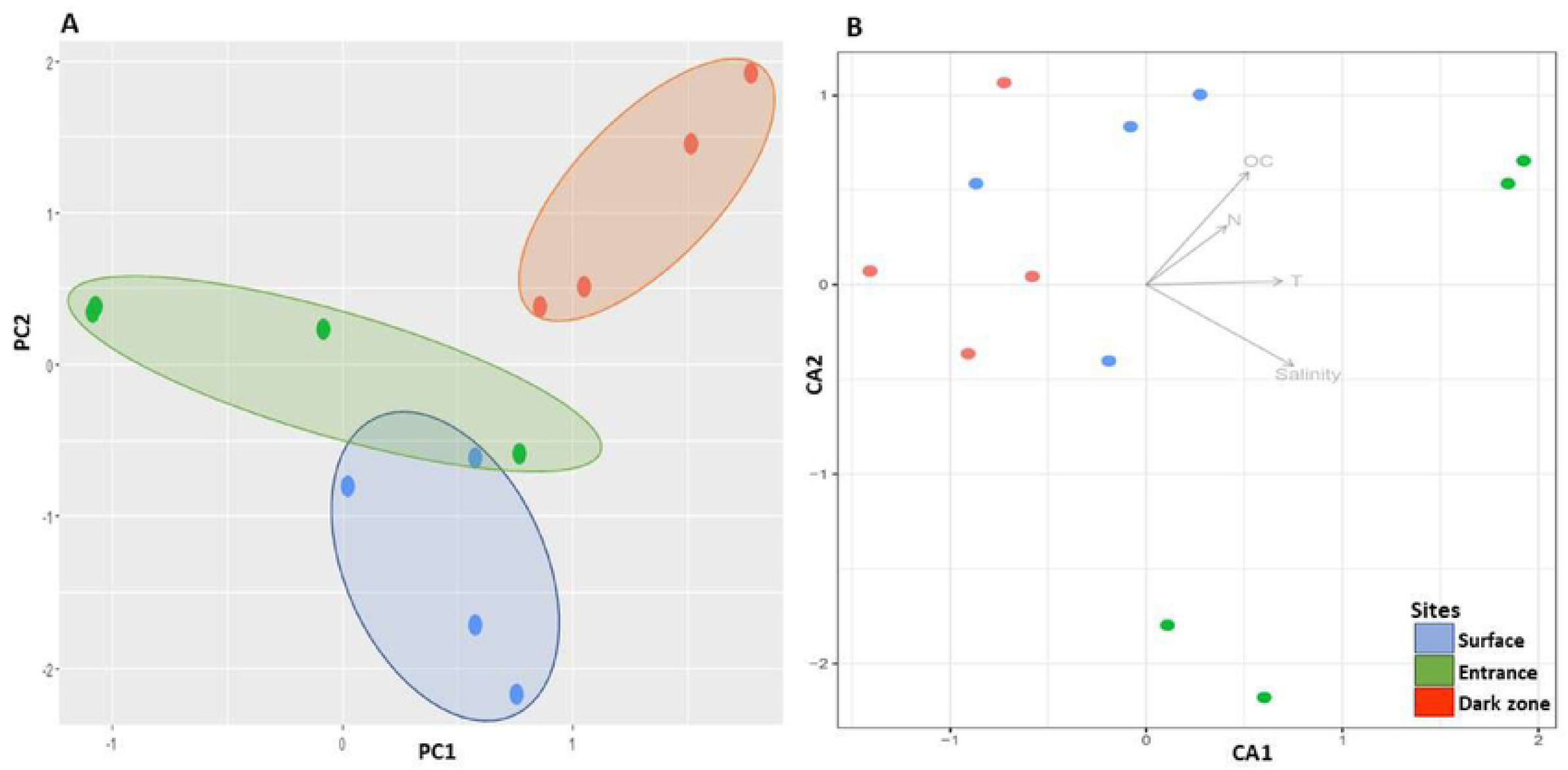
(A) Principal Coordinate Analysis (PCoA) plot based on weighted Unifrac results and (B) Canonical Correspondence Analysis (CCA) of microbiome data and environmental factors among the whole cave ecosystem, using Bray-Curtis distance and 999 permutations. Only environmental factors with p-values < 0.05 are marked at the graph. Samples from surface, entrance and dark zones are highlighted in blue, green, and orange, respectively.

**Fig 5.**
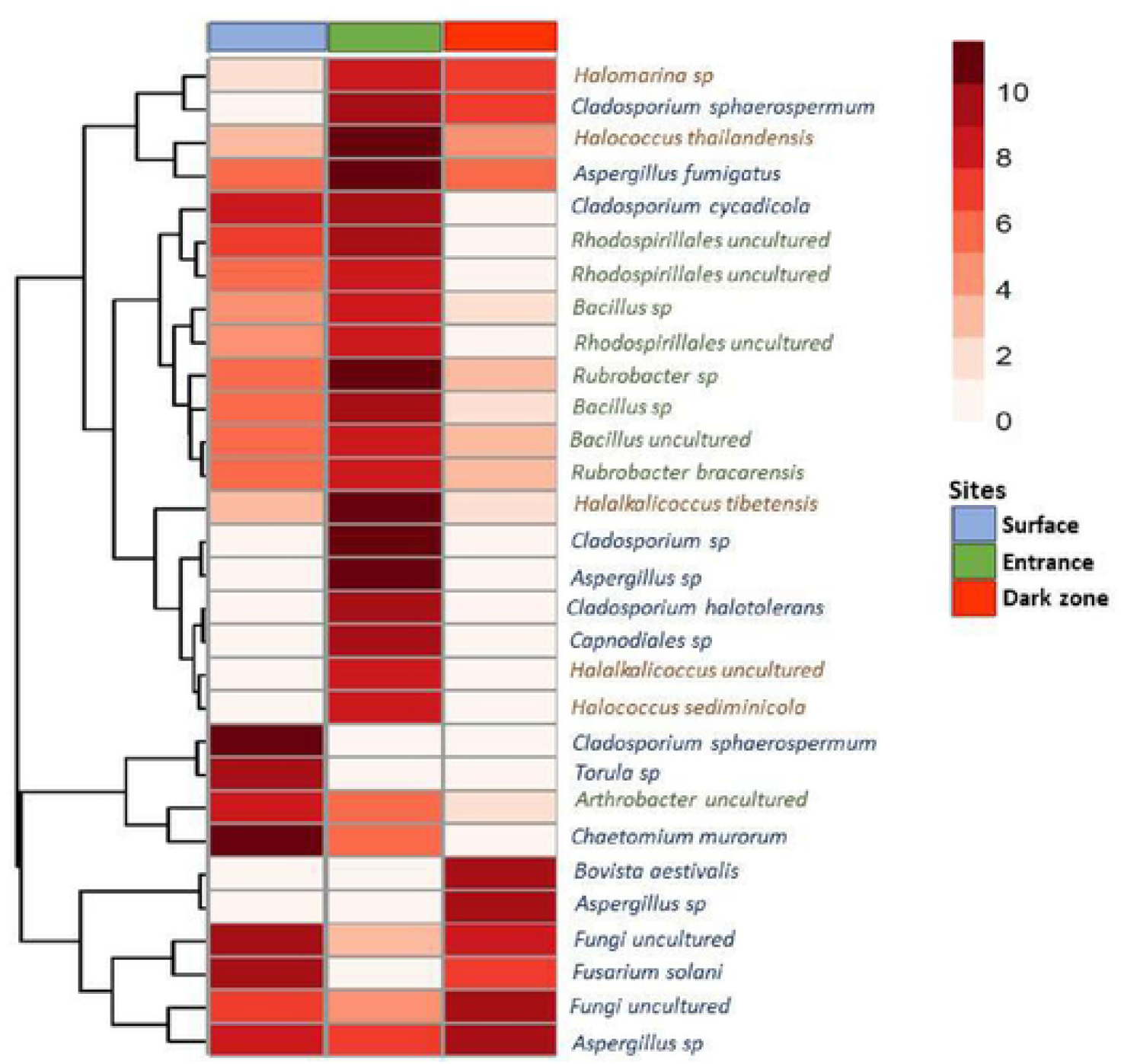
Heatmap of microbiome taxons with relative abundance ≥ 1% (core microbiome) from different habitats in the TR cave. Samples from surface, entrance and dark zones are highlighted in blue, green, and red, respectively. Fungal taxa are highlighted in dark blue, while bacteria are grey, and archaea are purple.

Canonical Correspondence Analysis (CCA) was used to identify possible relationships between microbial assemblages and local physicochemical variables at sampling sites (Fig 4B). The analysis of variance using distance matrices was used to find the best set of environmental variables that describe the community structure for each site sampled (p < 0.05). Among the tested variables, nitrogen (F2 = 1.96, p = 0.01), temperature (F2 = 1.49, p = 0.01), organic carbon (F2 = 2.02, p = 0.01), and salinity (F2 = 1.34, p = 0.02) were the main environmental drivers affecting the distribution of microorganisms. The variables showed a positive correlation with microbiomes at the cave entrance, while the surface and dark zone sites showed an opposite trend. These results clearly show a trophic and salinity gradient affecting the microbial structures at the cave entrance in relation to the other habitats.

## 4 DISCUSSION

Caves are widely considered as extreme oligotrophic environments (<5.0 mg C L^-1^), although this is largely based on research results on temperate cave environments. However, this assumption of oligotrophic conditions only holds true for subterranean environments that are relatively isolated from the surface, thus having very limited organic input [43]. The TR cave entrance and dark zone sites are relatively well connected with the surface environment, which is a rich source of organic matter as it is characterized by dense vegetation cover. This is reflected in the concentrations of organic carbon, which seems to accumulate at the cave entrance, with its concentration at dark zone site high, in the range of 100 mg g^-1^. This supports the conclusions of recent research that although both temperate and tropical cave communities are supported by organic inputs from allochthonous sources, and both types of systems show a trophic gradient from surface to dark zones, the tropical subterranean environments are not limited in energy input to the same degree as their temperate counterparts, due to the diversity and abundance of vegetation and its presence throughout the whole year [3, 5]. The environmental differences between tropical and temperate caves directly influenced the microbial communities within subterranean system. Tropical caves are a particularly understudied environment, and most of what we know about microbial communities was based on studies using culture-dependent methods [44 – 47]. While there are some published results generated by culture-independent approaches [6, 48], this is the first comprehensive study reporting the microbiome composition from a tropical cave, including prokaryotic and fungal assemblages through next generation sequencing.

TR cave sites hosted a heterotrophic microbiome and most of the dominant phyla identified in this study were previously also found in other cave systems worldwide (summarized in [2]). *Actinobacteria* and *Sordariomycetes* were the dominant classes at the dark zone site, which seems to be a general trend in limestone caves [4, 49]. *Actinobacteria* are known as typical heterotrophs, that play a crucial role in the carbon cycling through organic matter decomposition [50], and good competitors due to their wide metabolic capabilities which enable them to grow on organic compounds of varying complexity and the ability to produce several secondary metabolites, such as antibiotics [51, 52]. *Sordariomycetes* (Ascomycota), although their diversity remains underestimated, represent a recurrent fungal class in caves which also likely play an important role in the decomposition of organic matter, with their biomass (spores and mycelia) as an essential food source for many cave arthropod communities, such as isopods, collembolans, and protozoa [19, 20]. These saprophytic fungi show an aggressive colonization strategy, capable of surviving in extreme environmental conditions, and some fungi of this class, such as *Penicillium* and *Aspergillus*, are able to degrade rocks and solubilize minerals [53]. All these features regarding to cave microbiome are an important advantage to survive in low carbon and nutrient environments, as in dark zone at TR cave. Fungi are often studied in caves because to the concern in detecting pathogenic fungi. *Histoplasma capsulatumm*, the main pathogenic fungus in caves, was not found in our results. However, caves can present opportunistic fungi, which can be pathogenic under specific conditions, such as high concentrations of spores and dry air. Fungi species responsible for opportunistic infections have increased in recent decades [54]. *Aspergillus fumigatus*, for instance, was found in all sampled habitats and high abundance in entrance at the TR cave. *Aspergillus fumigatus*, an opportunistic fungus, is the main responsible for aspergillosis and it is necessary to inhale a high load of spores to develop the disease. Spores inhaled by healthy people are eliminated quickly by the immune system or just a weak allergy is developed. On the other hand, people with compromised immune systems can develop a serious pulmonary infection by inhaling a large load of spores [55]. Periodic monitoring of spore loads in TR cave is highly recommended in order to prevent people from entering the cave during periods with a high concentration of spores in the air, especially in drier periods. The use of cave for tourism and scientific purposes with no proper monitoring can lead to health problems for tourists, tour guides and researchers.

The chemical composition of the sediment in the TR cave shows the presence of cosmotropic and chaotropic salts. Cosmotropic salts are stabilizing salts, such as NaCl and KCl, that stabilize proteins, whereas chaotropic salts are recognized as destabilizing salts like MgCl_2_ that unfold proteins and increase solubility of hydrophobic chemicals [56], which makes the environment inhospitable for many microorganisms. Thus, the salinity can affect microbial communities through the osmotic effect and specific ion effects [57]. The salinity was found to be significantly higher at the cave entrance. Besides being a subterranean biodiversity hotspot, TR cave also has a cultural and religious value for the population from cities around the State Park of Terra Ronca. Since 1920, an annual religious festival attracts thousands of people to the cave entrance in August (dry season), and during this period the soil and sediment are highly compacted. In addition, fireworks containing residues rich in sulphur, potassium, and magnesium are used in large quantities [58]. Therefore, there are three factors possibly contributing to the increased salinity at this particular site: i) intense solar radiation at the cave entrance in comparison to the other two studied habitats and the consequent increase in water evaporation from the sediment; ii) sediment compaction during the religious festival decreases water retention and increases salt concentration in the substrate; and iii) intense and prolonged use of fireworks release salt-containing residues which accumulate in the sediment. All of these factors likely had an important effect on the microbiome composition and supported the dominance of halo-tolerant microorganisms at this site.

A significantly higher proportion of archaea, which surprisingly exceeded bacteria in relative abundance during the dry season, was found at the cave entrance site. The few research that has assessed archaeal diversity in caves emphasize the ecological importance of this group in subterranean environments due to their involvement in biogeochemical cycling, especially nitrogen and phosphorus [6, 8, 10]. Organisms belonging to the class *Halobacteria* (*Euryarchaeota*) were the dominant group here, being commonly found in environments with salinity higher than 3% and preference for neutral to alkaline pH [59], have an important role on nitrogen cycling – reducing nitrate and growing by denitrification [60], and hydrolyse insoluble phosphorus compounds to soluble compounds that can easily be assimilated by other organisms [61]. They are facultative phototrophic due the presence of an integral membrane protein known as bacteriorhodopsin [60, 62], a light-driven proton pump converting light energy into chemical energy, which may aid the growth under anoxic conditions, such as in a compacted sediment. All of these features highlighted the survival advantages of the *Halobacteria* at the cave entrance: presence of detectable inorganic phosphorus, under osmotic stress and anthropogenic disturbance.

Within the bacteria domain, *Bacilli* was the dominant class at the cave entrance, similarly to the prevalence reported from other cave systems, namely the Ozark cave, USA [51], and Brazilian caves [48]. Interestingly, in the Ararat Plain (Armenia), which represent a hydromorphic saline–alkaline soils, resembling those at the entrance of TR cave, the bacterial community was reported to be highly reduced, almost limited solely to *Bacilli*, which are able to remain viable and growing without competition from other bacteria [63]. Under such conditions, *Bacilli* show a specific survival strategy which includes the synthesis of special desiccation-resistant proteins, the accumulation of non-reducing sugars and the formation of dormant life stages (endospores). In terms of fungi, *Cladosporium halotolerans*, known as black fungi, also showed high prevalence at the cave entrance and this result is likewise consistent with the measured environmental conditions. This organism has been isolated from mine water in the Iron Quadrangle region (Minas Gerais, Brazil), is well adapted to harsh oligotrophic habitats on the surface and subsurface of rocks [64], high radiation, low water availability, long periods of desiccation, and shows high potential for removing Mn from the environment [65]. Thus, the microbiome at the cave entrance clearly shows the highest degree of microbial specialisation compared to the two other studied sites.

Microbiome composition and environmental parameters in TR cave clearly showed a distinction among the studied habitats. Canonical Correspondence Analysis (CCA) indicated organic carbon, nitrogen, temperature, and salinity as the main environmental drivers in the structure of microbial communities. Salinity was strongly related to the structure of the microbiome at the entrance cave, as previously discussed, and the others environmental drivers were related to dominant saprophytic microorganisms at surface and dark zone. Previous study at TR cave revealed the availability of carbon and nitrogen influenced the microbial strategies for organic matter decomposition and incorporation of those compounds into their biomass [5]. Now our results also support these environmental factors also influence the composition and structure of the microbial communities. The entrance of TR cave can be considered as an ecotone, the transitional zone between adjacent ecological systems (surface and dark zone), where the environment rapidly shifts from one type to another based on abiotic and/or biotic features. Even though many researchers consider ecotone an area with greater richness and diversity than each one of the systems, an ecotone can also support lower diversity if resources vary widely within it or if it is in an area under the influence of severe disturbances [66]. The anthropogenic impact and the unique habitat conditions, such high salinity and solar radiation, at the entrance of TR cave can promote the development of high endemism and dominance of few species, as already seen for invertebrate community in caves [67].

In summary, this study is the first to assess the microbiome structure in different habitats of a tropical cave system using high-throughput amplicon sequencing. The microbiomes at the surface and dark zone are composed mainly of heterotrophs microorganisms. This composition together with the relatively high organic carbon concentrations indicate the presence of a trophic network based almost entirely on detritivory. The influence of carbon and nitrogen, as seen in previous studies in TR cave, along with temperature, highlights those as the main drivers on the decomposing microorganisms, especially in the dark zone. Our study also shows for the first time the dominance of Haloarchea in a limestone cave, which may have an important ecological role in this environment as a phototrophic, phosphate solubilizing archaea, and nitrogen cycle players. Furthermore, these results show that anthropogenic changes can have profound implications for cave soil composition, microbiome structure, and, hence ecosystem functioning, should be considered in the future studies, alongside the commonly researched effects on micro- and macroinvertebrates and vertebrates.

## 5 Acknowledgments

This study was financed in part by the Coordenação de Aperfeiçoamento de Pessoal de Nível Superior - Brasil (CAPES) - Finance Code 001, Fundação de Amparo à Pesquisa do Estado de São Paulo (FAPESP, 2015/24763-9) for funding and supporting the project. We also thank Ramiro Hilário dos Santos, Jonas Eduardo Gallão, Maria José Rosendo da Costa and all member from the Laboratory of Subterranean Studies for assisting in the field sampling. We appreciate the Grupo Bambuí de Pesquisas Espeleológicas (GBPE) for use permission of the maps and the environmental agencies for the permission to collect: Biodiversity Authorization and Information System / Chico Mendes Institute for Biodiversity Conservation (SISBIO / ICMBIO); Goiás - Secretary of Environment, Water Resources, Infrastructure, Cities and Metropolitan Affairs (SECIMA). The authors are grateful to the Postgraduate Program in Ecology and Natural Resources (PPGERN/UFSCar) for the infrastructure. We thank Angélica Maria Penteado Martins Dias and Luciana Bueno dos Reis Fernandes (INCT-HYMPAR/UFSCar) for the chemical analysis of the substrates by scanning electron microscopy (SEM) coupled to energy dispersive spectroscopy (EDS).

## 7 Supporting information

**Fig S1.** Detailed map of the Terra Ronca - Malhada subterranean system. Highlight for the (A) Terra Ronca I (TR cave) and (B) Terra Ronca II caves. Map: Bambuí Speleological Research Group – GBPE.

**S2 Appendix.** Prokaryotical OTU (Operational Taxonomic Unit) table.

**S3 Appendix.** Fungal OTU (Operational Taxonomic Unit) table.

## 10 Data Availability

Raw sequence data from this study can be downloaded from National Center for Biotechnology Information (NCBI) Sequence Read Archive (SRA) with accession number PRJNA723998 (16S) and PRJNA724003 (ITS).

## 11 Conflict of Interest

The authors declare that they have no conflict of interest.

## 12 Author Contributions

PAULA, CCP conceived and designed the study, responsible for chemical and molecular analyzes and wrote the paper with input from all authors. SELEGHIM, MER was in charge of the funding budgets and wrote the paper. BICHUETTE, ME contributed to data analyses, coordinated the field sampling program and contributed to writing of the paper. SIROVÁ, D and SARMENTO, H contributed to data analyses and writing of the paper. FERNANDES, CC and KISHI, LT were involved in the sequencing of the 16S samples.

## 13 Funding

The study was supported by Coordenação de Aperfeiçoamento de Pessoal de Nível Superior - Brasil (CAPES) - Finance Code 001 awarded to CCPP and Fundação de Amparo à Pesquisa do Estado de São Paulo (FAPESP, 2015/24763-9) awarded to MHRS.

## Notes

### Competing Interest Statement

The authors have declared no competing interest.

